# Scalable single-cell isoform profiling with sequencing-by-expansion

**DOI:** 10.64898/2026.07.15.738809

**Authors:** Christophe H. Georgescu, Ghamdan Al-Eryani, Allison Brookhart, Jagdeesh Chandrasekar, Stephanie J. Yaung, Megan Rogers-Peckham, Megan Freer, Michael E. Kartje, Houlin Yu, Akanksha Khorgade, Charlotte Yang, Lacey McGee, Kendall Berg, Cynthia Cech, Salka Barrett, Anasha Arryman, Dan A. Bartlett, Adam Slamin, Sophie Low, Dan Dubinsky, Michelle Cipicchio, Nir Hacohen, Taylor Lehmann, Niall J. Lennon, Victoria Popic, Chen Zhao, Marc Prindle, John Mannion, Melud Nabavi, Brian J. Haas, Mark Kokoris, Aziz M. Al’Khafaji

## Abstract

Single-cell RNA sequencing has transformed our understanding of cellular systems, yet the reliance on short-read sequencing restricts analysis to gene-level quantification and obscures the immense biological diversity generated by alternative splicing. While long-read sequencing technologies can capture full-length RNA and resolve transcript isoforms, current platforms remain constrained by throughput and high per-base costs, rendering them impractical for modern million-cell applications. To address this critical limitation, we developed and optimized sequencing-by-expansion (SBX) chemistry for high-throughput single-cell RNA isoform profiling. Integrated within the AXELIOS 1 sequencing platform, SBX employs a unique biochemical conversion process that transforms complementary DNA into expanded surrogate high signal-to-noise polymers called Xpandomers which are sequenced via translocation through a dense nanopore array yielding over 9.5 billion reads in a two-hour run. To leverage this unique data type for long-read single-cell RNA isoform sequencing, we developed the Consensus UMI Deduplication using Longest Length (CUDLL) algorithm, which computationally consolidates variable-length raw SBX reads into single, high-fidelity consensus reads, elevating sequence accuracy to 99.83% and maximizing per transcript read length. We demonstrate that this consensus approach successfully captures the vast isoform diversity of single-cell libraries and enables the accurate measure of differential isoform expression across distinct cell types in peripheral blood mononuclear cells. Furthermore, SBX coupled with CUDLL efficiently resolves T-cell and B-cell receptor clonotypes directly from whole-transcriptome libraries without the need for VDJ-specific target enrichment. Ultimately, this work establishes SBX and the AXELIOS 1 as a transformative platform for high-scale single-cell isoform sequencing.

## Introduction

The advent of single-cell RNA sequencing has revolutionized our understanding of cell biology, providing unprecedented resolution into the diversity of gene expression programs and biological mechanisms underlying health and disease. Despite the wealth of discoveries made over the past decade driven by advances in transcriptomics, reliance on gene-level expression quantification has masked the true diversity of transcripts arising from alternative splicing. Alternative splicing is a core regulatory process that orchestrates the differential inclusion or exclusion of exons during mRNA maturation, ultimately modulating the final transcript structure, which contains regulatory motifs and protein coding information ^1–3^. Transcript isoforms arising from the same gene can encode proteins with slightly altered, markedly different, or even opposing functions ^4^. This functional variation constrains the utility of gene-level expression measurements alone. To address this limitation, long-read approaches have been recently leveraged to capture full-length RNA data, enabling isoform expression analysis from single-cell and spatial samples ^5–9^. While this transformative paradigm enables the bridging of systems, molecular, and structural biology, existing methodologies remain unable to scale to modern single-cell throughputs of millions to billions of cells.

While short-read platforms such as Illumina remain the standard for scRNA-seq due to their throughput and cost-effectiveness ^10^, their limited read length hinders the direct inference of isoform structures or allele-specific expression ^11–14^. Long-read sequencers, such as Oxford Nanopore Technologies (ONT) or Pacific Biosciences (PacBio), circumvent these shortcomings by generating reads that span full or near-full transcript lengths ^5,13,15–17^. Consequently, emerging long-read single-cell workflows now enable detailed transcriptomic analyses, including the detection of RNA isoforms, fusion transcripts, RNA edits, allele-specific expression, and splicing quantitative trait loci ^14,18–21^. However, long-read scRNA-seq confronts a critical tradeoff, higher-throughput platforms like ONT yield high read counts but exhibit elevated error rates ^8,22^, whereas PacBio HiFi offers higher accuracy (<1% error rate) but at the cost of reduced throughput and higher expense ^23^. Recent innovations have sought to mitigate these constraints. For example, MAS-seq (commercially Kinnex) leverages programmatic cDNA concatenation to increase throughput on PacBio instruments by 16-fold ^16^. Despite these advances, achieving a practical balance of accuracy, throughput, and cost for routine, million-cell scale single-cell isoform profiling remains a significant challenge.

Here we introduce and evaluate sequencing-by-expansion (SBX) for highly scalable single-cell isoform sequencing. SBX employs a biochemical conversion process that transforms cDNA into expanded surrogate polymers called Xpandomers ^24^. This expansion is achieved through the incorporation of specialized expandable nucleotide triphosphates (X-NTPs) during enzymatic synthesis, followed by a cleavage step that releases the tethered backbone, generating a high signal-to-noise extended polymer approximately 50-fold longer than the input DNA. These Xpandomers are subsequently translocated through nanopores integrated within a high-density CMOS-based sensor array enabling massively parallel single-molecule sequencing ^24^. The pairing of high signal-to-noise reads with rapid parallel translocation rates yields immense sequencing throughput of approximately 4.8 billion reads per hour on the AXELIOS 1 platform. Together, these features position SBX as a scalable platform for rapid, high-capacity sequencing, with utility ranging from time-sensitive applications such as rapid diagnostics to large cohort sequencing ^25^. The unique sequencing throughput and read lengths of SBX are particularly compelling for enabling high-scale single-cell RNA isoform sequencing. We demonstrate the capacity for SBX to efficiently capture transcript isoforms from peripheral blood mononuclear cells (PBMCs) using the 10x Genomics 5’ workflow. To maximize read accuracy and length, we developed a specialized workflow and consensus algorithm for SBX reads, which greatly increases read utilization and isoform identification. We observed high levels of concordance between the AXELIOS 1 and Revio platforms, with enhanced capture for lowly expressed or longer transcript isoforms respectively. Finally, we demonstrate SBX’s ability to efficiently capture BCR and TCR transcripts, underscoring a key capability to integrate key immune clonotype information.

## Results

To evaluate the utility of SBX for single-cell RNA isoform studies, we benchmarked AXELIOS 1 against PacBio Kinnex, a state-of-the-art platform for long-read RNA isoform sequencing. We used PBMC samples captured with the 10x Genomics 5’ v2 chemistry to assess both base-level accuracy and isoform-level agreement between the two platforms (**Fig. 1A**). To manage the massive scale of the SBX data (∼9.5B reads generated in a 2h sequencing run) and address its inherent error rates, we developed a new single-cell workflow designed to maximize both computational efficiency and data accuracy (**Fig. 1B**). Importantly, we tailored this workflow to specifically model the unique biochemical dynamics of Xpandomer generation, processing, and sequencing, which lead to variable read termination positions with respect to the parent template molecule. To that end, we developed the **C**onsensus **U**MI **D**eduplication using **L**ongest **L**ength (**CUDLL**) algorithm, which consolidates variable-length raw reads into their accurate, singular molecular representations. Within each UMI group, CUDLL selects the longest read as the anchor (maximizing transcript coverage) and collapses the remaining reads into a single representative CUDLL sequence **(Fig. 1C)**.

**Fig. 1:**
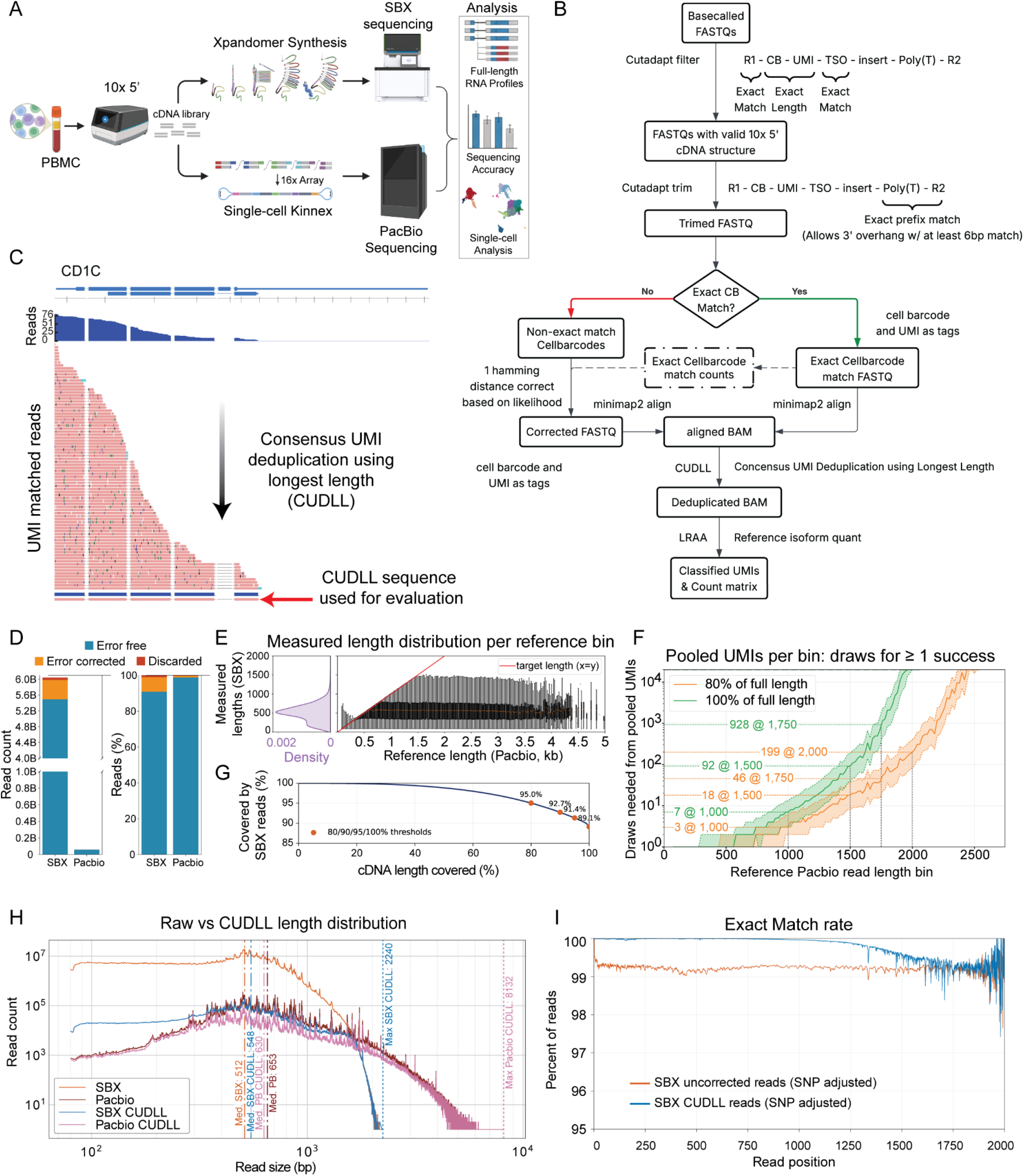
Overview of evaluation framework, data processing, and primary metrics. **A)** Evaluation workflow. **B)** Diagram of the data processing pipeline, starting from raw reads to CUDLLs and expression matrices. **C**) Example alignment of raw SBX reads derived from the same UMI and the resulting CUDLL. Introns are excluded to better visualize annotated exons. **D**) Absolute read count and percentage of adapter filtered reads that are either an exact match to a whitelist barcode, error-corrected to a whitelist barcode, or are discarded. **E**) Left panel, adapter trimmed SBX Read length distribution (purple). Right panel, adapter trimmed SBX Read length distribution according to the longest PacBio read for the same CB-UMI, grouped by 25 bp length bins. Boxplots indicate 0.01, 0.25, 0.5, 0.75, 0.99 SBX length quantiles. **F**) Expected number of copies of a CB-UMI needed for at least one to be 80% (orange) or 100% (green) of the full cDNA length at different target likelihoods (quartiles). **G)** Fraction of cDNA molecules covered by SBX reads at a given target length. **H**) Aligned read length distribution of filtered reads and CUDLLs for both SBX and PacBio. **I**) Exact match rate per position along the read for SBX filtered reads and CUDLLs, filtering out VDJ, HLA and MT regions, and only using CB-UMI that are FSM/ISM/se_FM/se_IM in PacBio. PacBio based DeepVariant VCF and UMI matched errors shared with PacBio are considered true polymorphism or PCR errors and counted as matches.

To quantify the fraction of high-confidence reads, we first computed the number of raw sequences, which have a valid cDNA structure and correct cell barcode. We found that 63.37% of raw SBX reads (6.04B of 9.54B) contained the correct 10x 5’ cDNA adapter structure (with an exact TSO match and expected CB and UMI sequence length) as compared to 86.17% of PacBio Kinnex S-reads (60.07M of 69.7M) (**Table S1**). For SBX, 90.88% of these retained reads perfectly matched the whitelist cell barcode, while 8.12% had a Hamming distance of 1 (a single-nucleotide mismatch) and were corrected. This resulted in 99% of SBX reads with the correct 10x 5’ structure being usable **(Fig. 1D)**. By comparison, 98.86% of PacBio barcodes perfectly matched the whitelist and 0.97% were correctable, yielding a 99.83% usable read rate. Ultimately, we obtained a median of 1,414,508 reads/cell for SBX and 14,939 reads/cell for PacBio (**Table S1**). Following CUDLL deduplication, this generated 2,134 genes/cell (8,814 UMIs per cell) for SBX and 1,067 genes/cell (3,804 UMIs per cell) for PacBio.

Next, we evaluated the read length distribution of SBX and the structural improvements conferred by CUDLL. To distinguish premature Xpandomer read termination from cDNA truncation caused by RNA degradation, incomplete reverse transcription, or PCR artifacts, we compared SBX and PacBio reads with length greater than 1,500 bp that shared the same cell barcode and UMI (CB-UMI). We found a median SBX read length of 551 bp (75th percentile: 726 bp), indicating the current upper limits of SBX read lengths. By calculating sampling probabilities and expected read depths **(Fig. 1F)**, we found that capturing 100% of a transcript requires exponentially more reads for molecules >1,500 bp. Conversely, 80% coverage is reliably achieved with significantly fewer reads, even for transcripts spanning 1,000 to 2,000 bp (**Fig. 1F**). We observed 89.1% of cDNAs to be sequenced end-to-end by SBX, underscoring the rate of complete cDNA coverage from 10x single-cell libraries. (**Fig. 1G**). Applying CUDLL increased the SBX read length distribution from a median of 512 bp to 548 bp, while running CUDLL on PacBio Kinnex reads resulted in a decreased median read length from 653 bp to 630 bp. This result is due to the combination of PacBio’s end-to-end sequencing and the presence of a population of single-read per UMI cDNAs below 400 bp which lowers the median size distribution of reads post-deduplication. When comparing the maximum observed read lengths, SBX reads reached 2,240 bp, whereas PacBio captured lengths up to 8,132 bp, reflecting the current capacity of each platform to unambiguously identify long transcripts **(Fig. 1H)**.

To further assess primary read-level metrics, we quantified the sequence level accuracy of the SBX platform in the context of single-cell cDNA sequencing. As apparent errors can arise from natural variation, reverse transcription errors, or early PCR errors rather than from Xpandomer synthesis or SBX sequencing, we filtered apparent errors using Pacbio-only VCF calls, then further using differences shared between PacBio and SBX reads with the same CB-UMI (see **Methods**). Specifically, a mismatch was counted as a platform-specific error only when it was observed uniquely in SBX or PacBio, but not in both datasets (**Fig. 1I**). Across shared-cell CUDLLs, the VCF-adjusted per-base match rate was 99.53% for SBX CUDLL reads, and 99.92% for PacBio CUDLL reads, with errors primarily arising from substitutions **(Table S2)**. In comparison, we found a per-base match rate of 99.01% for SBX, and 99.84% for PacBio in uncorrected reads. When filtering for SBX CUDLLs formed from at least 3 reads, the overall per-base match rate increased to 99.83%, and further increasing to 99.86% for CUDLLs formed from 10 reads (**Table S2**). To correct for biological or PCR derived variation, we additionally filtered out errors observed on both sequencing platforms, resulting in accuracy rates of 99.96% for SBX and 99.97% for PacBio (**Table S2**). Notably, sequence fidelity decreased significantly toward the 3’ end of the read, reflecting a drop in the read coverage necessary for CUDLL to perform consensus corrections **(Fig. 1I)**.

To explore SBX’s capacity for isoform profiling from single-cell libraries, we leveraged LRAA to classify CUDLL reads according to the SQANTI isoforms structure classification framework as well as quantify reference annotated gene and isoform expression ^26,27^. While PacBio generated proportionally more full-splice matches (FSMs) and fewer incomplete splice matches (ISMs) compared to SBX, the immense throughput of SBX ultimately yielded over 50% more total FSM reads (**Fig. 2A-B)**. Of the distinct isoforms with identified FSM assignments, 69.4% were shared between the two platforms, while 21.2% were unique to SBX, and 9.4% were unique to PacBio. Further, as expected, the FSMs unique to PacBio corresponded to cDNAs from longer transcript, which likely featured splice junctions at the distal ends of the molecules that SBX failed to capture (**Fig. 2C**).

**Figure 2:**
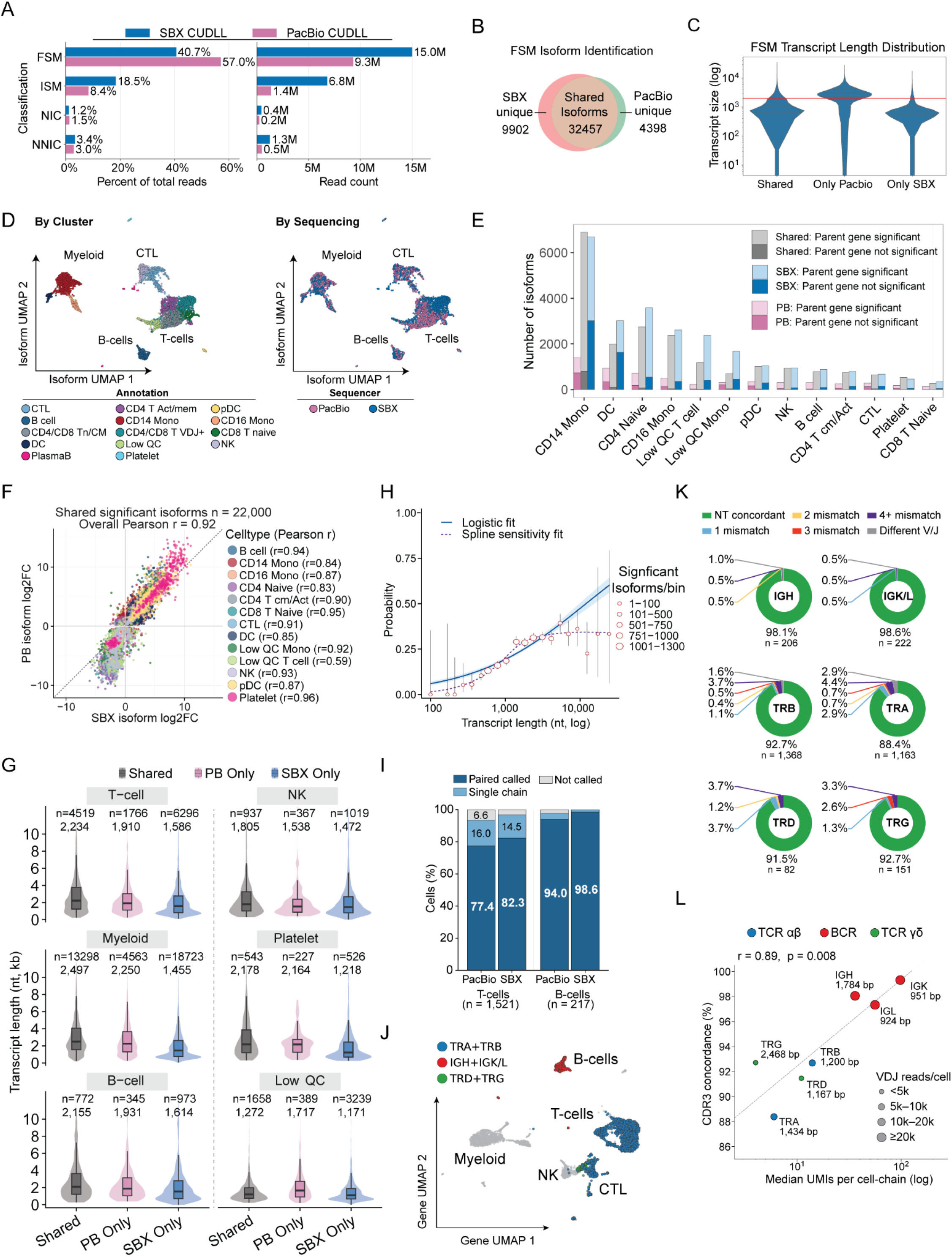
SBX efficiently resolves ∼1.5 kb isoforms and antigen-receptor sequences for single-cell data. **A**) LRAA-SQANTI classification for SBX and PacBio CUDLLs, shown as (left) the percentage of total and (Right) as absolute counts. FSM: Full Splice Match, ISM: Incomplete Splice Match, NIC: Novel In Catalog, NNIC: Novel Not In Catalog. **B**) Overlap of FSM isoforms detected by SBX and PacBio, identifying 32,457 shared isoforms, 9,902 SBX unique isoforms and 4,398 PacBio unique isoforms. **C**) Reference transcript length distributions for shared, PacBio only and SBX only FSM. The red bar indicates 2 kb. **D**) PacBio and SBX harmony Integrated UMAP based on isoform expression, with single cells colored by cell-type annotation (left) or by sequencing platform (right). **E**) Cell-type resolved counts of significant isoforms stratified as shared, SBX-only or PacBio-only and further annotated by whether the corresponding parent gene is significant at the gene-expression level. **F**) Concordance of isoform log2 fold-changes for 22,000 isoforms significant in both platforms, with points colored by cell type (overall Pearson correlation of r = 0.92 between SBX and PacBio). **G**) Transcript-length distributions for shared, PacBio-only and SBX-only significant isoforms across major lineages. **H**) Logistic model estimating the probability that a non-shared significant isoform is PacBio-only as a function of transcript length. Points show binned observations, point size denotes the number of significant isoforms per bin, and the spline fit provides a sensitivity analysis for the length-dependent trend. **I**) Fraction of T cells and B cells with paired, single-chain or no V(D)J calls in PacBio and SBX. **J**) Gene expression UMAP overlaid with filtered V(D)J calls, showing localization of TRA+TRB, IGH+IGK/L and TRD+TRG calls. **K**) Nucleotide-level concordance of matched V(D)J chain calls between SBX and PacBio, summarized by receptor chain. CDR3; complementarity-determining region 3; NT, nucleotide; UMI, unique molecular identifier; V/J, variable/joining gene usage. **L**) The relationship between SBX median UMI depth per cell-chain and PacBio to SBX CDR3 concordance across receptor chains. Dot color denotes receptor class and dot size denotes V(D)J reads per cell, showing that chain-level concordance increases with UMI support. The number next to the chain depicts the length of the aligned chain (as per GTF).

Having characterized the isoform discovery potential of each platform, we next sought to determine how these technical differences affect biological interpretation (**Fig. 2D-L; Table S3**). Across 2,647 same cell cluster matched barcode pairs, isoform-level testing (Wilcoxon rank sum test, Benjamini-Hochberg adjusted p-value = 0.05, log2FC ≥ 0.25) returned 62,509 significant isoform cell-type calls (≈53,189 unique isoforms), with 22,000 (35.2%) recovered by both platforms, 32,442 (51.9%) by SBX alone, and 8,067 (12.9%) by PacBio alone. At the gene level, this corresponded to 21,711 genes with at least one significantly differentially expressed isoform (of these genes, 12,120 were shared, 6,852 were SBX-only, and 2,739 were PB-only; **Fig. 2E**). Of the shared genes, 92.2% (11,169 / 12,120) also reached significance at the parent-gene RNA level, whereas a substantially larger fraction of platform unique calls (∼33% PB-only, ∼35% SBX-only) were significant at the isoform level only, consistent with these representing transcript-usage shifts that are masked at the gene-summary level.

Among isoforms identified as differentially expressed by both platforms, isoform fold changes were highly correlated across 12 of 13 cell types (Pearson r = 0.83 - 0.96), with Low QC T cells the lone outlier at r = 0.59 (**Fig. 2F**). Platform-unique calls were systematically shorter for SBX and longer for PacBio across all cell types (**Fig. 2G**). Longer transcripts were progressively more likely to be detected only by PacBio, with each 10-fold increase in transcript length raising the chance of PacBio only detection roughly 4-fold (**Fig. 2H**; logistic regression, p < 1 × 10^−71^).

A key objective at the intersection of immunology and single-cell technologies is the resolution of VDJ regions that define T- and B-cell receptor specificity and enable clonotype-resolved interpretation in addition to gene/isoform expression analysis of cellular states. Given the length and accuracy profile of SBX-CUDLL reads, we evaluated their capacity to accurately recover these immune transcripts directly from single-cell whole transcriptome libraries. To evaluate VDJ transcript recovery, we used PacBio VDJ calls as a reference and analyzed concordance across the matched set of 1,521 T cells and 217 B cells. To reduce the likelihood of assigning antigen-receptor sequences derived from ambient RNA, we first classified cells as TCR or BCR expressing cell types on the basis of gene-expression profiles (see **Methods**). SBX recovered at least one productive VDJ chain in 100% of B cells and 96.8% of T cells, compared with 97.7% and 93.4%, respectively using PacBio (**Fig. 2I-J**). Of these, productive paired-chain recovery reached 98.6% with SBX versus 94.0% with PacBio for B cells, and 82.3% versus 77.4% for T cells (**Fig. 2I**). To assess the sequence accuracy of SBX VDJ calls, we compared the top CDR3 nucleotide sequence assigned to each cell across both platforms, again utilizing PacBio calls as a reference. Among the 3,192 chain calls detected by both platforms, 91.9% exhibited exact nucleotide-level CDR3 sequence identity **(Fig. 2K)**. Chain-level concordance ranged from 88.4% (TRA) to 99.3% (IGK), while paired VDJ level concordance reached 82.1% for TRA + TRB and 97.1% for IGH + IGK/L **(Fig. 2K**). Furthermore, per-chain CDR3 concordance scaled monotonically with the median SBX UMI sampling depth per cell-chain (Pearson r = 0.89) (**Fig. 2L**), with BCR chains exhibiting a 3-15x higher per-chain UMI capture rate than TCR chains. While raising the UMI and read depth thresholds improved CDR3 concordance, it came at the expense of overall VDJ recovery, eliminating not only low-fidelity calls but also a substantial fraction of high-accuracy assignments. Together, these findings show that SBX-CUDLL reads can recover productive TCR and BCR clonotypes from single-cell whole transcriptome libraries without the need for VDJ specific cDNA enrichment, allowing scalable linking of antigen-receptor identity with isoform-resolved cellular phenotypes.

## Discussion

The diversity of isoforms underlying the transcriptome has long been obscured in single-cell studies due to technical constraints of short-read sequencing. Our evaluation establishes SBX as a powerful sequencing approach capable of bridging the critical gap between >1 million cell single-cell applications and long-read sequencing to enable RNA isoform resolution at this scale. While the biochemical dynamics of Xpandomer generation introduce truncation in longer molecules, SBX overcomes this limitation through sheer throughput; two hours of sequencing yielded over six billion usable reads, with a median of ∼1.4 million reads/cell. This massive data volume confers the UMI redundancy necessary to enable our consensus-based CUDLL workflow, which outputs error-corrected and length-maximized deduplicated reads. This process enables accurate identification of splice-resolved isoform structures, even for transcripts exceeding 2,000 bp. Consequently, despite generating a higher per-read proportion of ISM, the sequencing depth of SBX ultimately captures a larger absolute number of FSM reads. While raw SBX reads exhibited an overall 99.0% match rate, leveraging CUDLL with a threshold of at least three supporting reads elevated the SNP-adjusted accuracy to 99.83%. Notably, the same depth-driven accuracy generalises to recombined immune-receptor loci, with SBX recovering productive VDJ chains in 100% of B-cells and 96.8% of T-cells at 91.9% nucleotide-exact CDR3 concordance with PacBio called CDR3 reads. However, the CUDLL read sequence fidelity decreases significantly toward the 3’ end of the reads, a consequence of the variable read lengths across the parent molecule. As reads terminate at different points, the depth of the read pileup decreases at the distal 3’ end, limiting the algorithm’s ability to perform robust consensus correction in those terminal regions.

While SBX successfully captures a vast diversity of transcript isoforms, its current sequence length constraints limit isoform discovery for longer transcripts. This defines a clear trajectory for future biochemical optimization, with enhancements in SBX read lengths allowing for greater capture across transcript length scales. Nonetheless, our coverage analysis indicates that its high-throughput sequencing reliably achieves complete coverage across 89.1% of single-cell cDNAs captured, which is sufficient for robust differential isoform and alternative splicing analyses. In this work, we demonstrate the utility of SBX sequencing paired with the CUDLL consensus workflow as a powerful sequencing solution for routine and high-scale isoform-resolved single-cell profiling. Such technologies are essential to catalyze the transition from gene to isoform level transcriptomics, a paradigm which will bridge systems, molecular, and structural biology and drive waves of discovery across the basic and translational sciences.

## Methods

### SBX library preparation and sequencing

10× Genomics Single Cell 5’ v2 full-length cDNA libraries generated from PBMC (AllCells; Lot # 3082436 25/Apr 22 with 10 million cells) that were generated and evaluated previously by Webber et al. ^13^ was processed for SBX workflow. PCR was used to introduce SBX specific sequence handles. PCR amplification was performed using 2 ng cDNA in a 50 μL reaction using KAPA HiFi HotStart ReadyMix, with 1 μM of each primer for 10 cycles (64°C annealing; 72°C extension for 1 min). The adapted cDNA product was purified with KAPA HyperPure beads at a 0.6x ratio and quantified on a Qubit 4 Fluorometer (Thermo Fisher Scientific, Waltham, MA) using the dsDNA HS Assay Kit. Subsequently, 50 ng of the PCR-amplified library was subjected to linear amplification for 6 hours followed by a second KAPA HyperPure bead purification at a 2x ratio. Final library quantification was performed on the Qubit 4 using a custom-built assay (Roche Sequencing Solutions, Seattle WA).

Sequencing by Expansion (SBX) is performed in two steps: Xpandomer (Xp) synthesis and sequencing measurement of the Xp. First, SBX adapted DNA libraries were loaded into an AXELIOS 1 prototype synthesis instrument to serve as templates for encoding into Xpandomers, which act as highly measurable surrogate molecules. These encoded Xp constructs were then sequenced for 2 hours on the AXELIOS 1 prototype sequencing instrument. The instrument’s 8M sensor array enables highly parallelized measurement of these surrogate molecules resulting in exceptionally high throughput. Finally, real-time basecalling was performed directly on the instrument, with the output subsequently converted into FASTQ format.

### SBX reads processing

Raw sequencing data in FASTQ format were processed to identify and isolate the 10x Genomics Chromium Single Cell 5’ v2 library structure. Due to the distinct error profile of SBX sequencing, which inherently produces insertions alongside substitutions, we implemented a stringent filtering approach using Cutadapt ^28^. The 5’ adapter structure required an exact match comprising 12 bp of the R1 PCR handle, 26 random nucleotides (16 bp cell barcode and 10 bp UMI), and the 13 bp Template Switching Oligo (TSO). Because Xpandomer synthesis exhibits variable template termination, 3’ adapter trimming was relaxed to match and trim a minimum 6 bp match to the RT adapter, with terminal poly-A tracts removed regardless of length.

Cell barcodes (16 bp) and UMIs (10 bp) were extracted from their now expected positions using a custom tool and evaluated against the standard 10x 5’ v2 whitelist. Reads with exact barcode matches were retained and output as adapter-trimmed, tagged FASTQs. To maximize data retention, reads with unmatched barcodes underwent error-correction at a Hamming distance of 1, utilizing posterior probability distributions analogous to the Cell Ranger pipeline ^29^. Reads that could not be corrected or did not meet the minimum length threshold were discarded. All processing steps were integrated into a configurable pipeline to enable automation and parallelization.

### PacBio reads processing

To minimize differences attributable to upstream processing, PacBio S-reads were first processed through Skera and Marti ^13^, reads were then reoriented to the forward strand based on adapter orientation and subsequently processed through the same pipeline described above for SBX reads.

### Alignment

Reads were aligned to the human reference genome (refdata-gex-GRCh38-2020-A, 10x Genomics) using Minimap2 ^30 31^. SBX reads were aligned with the options “-ax splice -uf -G 1250k --eqx”, while PacBio reads were aligned using “-ax splice:hq -uf -G 1250k --eqx”. Annotated splice junctions were provided to Minimap2 via the “--junc-bed” parameter to improve spliced alignment accuracy.

### CUDLL Deduplication

Given the unique length characteristics of Xpandomers and the throughput of SBX sequencing, we developed CUDLL (Consensus UMI Deduplication usingLongest Length), a two-pass deduplication tool that collapses reads sharing a cell barcode and UMI while generating a consensus sequence prioritizing read length.

#### First pass

Starting from a coordinate-sorted BAM, overlapping reads are first clustered by cell barcode, then by UMI and strand. Within each cell barcode, UMI clusters are processed in descending order of read support. Each UMI is compared against higher-support UMIs, and clusters within a Hamming distance of 1 are merged into the larger group. Once all clusters are finalized, splice junction patterns are evaluated per read, and the pattern with the greatest support (with compatible subsets counted as supporting evidence) is selected as the representative isoform. Among reads matching the selected splice pattern, the read with the greatest number of aligned bases is chosen as the consensus template, defining alignment boundaries and splice junctions. A consensus sequence is then built using all reads in the cluster as supporting evidence, leveraging CIGAR strings where possible. At ambiguous positions, the reference base (if a reference is provided or CIGAR “=“ and “X” are used) is preferred unless more than half of the reads at that position support a different base; when two non-reference bases compete, majority support is used, and in case of tied support, base quality scores are used to arbitrate. The resulting consensus read (CUDLL) retains the read ID of the template read and carries two custom tags: “DC” (total read support) and “UD” (UMI sequences of 1-mismatch UMIs merged). Most original per-read tags are stripped, as they may no longer be valid for the consensus sequence; however, the “nm” tag (number of mismatches) is recomputed if a reference FASTA is provided. The output BAM is then sorted by cell barcode in preparation for the second pass.

#### Second pass

The second pass resolves UMIs that may have mapped to multiple genomic loci due to alignment ambiguity, which can arise from short Xpandomers or paralogous sequences. CUDLLs are processed per cell barcode in descending order of read support (“DC” tag). UMIs that are identical or within a Hamming distance of 1 are flagged as candidate duplicates and subjected to pairwise sequence alignment; pairs with ≥95% sequence identity are merged. Unlike the first pass, the representative CUDLL in the second pass is selected based on aligned length rather than read support, as longer alignments should provide less ambiguous locus assignments. The “DC” and “UD” tags are updated accordingly upon merging.

This two-pass strategy has several important properties. First, true UMI collisions in different loci are generally preserved as separate entries due to the sequence identity threshold. Second, fusion transcripts generating supplementary alignments are retained as two alignments under the same cell barcode and UMI. Third, read-through molecules, which should carry a second UMI, behave like fusions and are preserved as two CUDLLs under the same cell barcode and UMI of the first read (automated detection of such events is planned for a future release). Fourth, 1-mismatch UMI copies mapping to alternative loci alone can still be correctly deduplicated, this unlike existing approaches such as Cell Ranger.

### Quantification and CUDLL Classification

Gene and isoform expression were quantified using LRAA v0.15.0 ^27^ in reference-based quantification-only mode with the basic single-cell pipeline. The “HiFi” parameter was set to false for SBX data and true for PacBio data. CUDLL classification was performed using LRAA’s integrated SQANTI-like read classification utility.

### SBX Read Length Distribution

PacBio read lengths were used as ground-truth estimates of insert size (excluding poly-A tails and adapters) for each cell barcode-UMI (CB-UMI) combination. For each CB-UMI, all corresponding SBX read lengths were collected. The dataset was then filtered to retain only CB-UMI pairs in which: (1) the cell barcode belonged to the shared set of high-confidence called cells, and (2) both SBX and PacBio reads were assigned to the same gene by LRAA, reducing confounding sources of variation. PacBio read lengths were binned in 25 bp increments, and the 1st, 25th, 50th, 75th, and 99th percentiles of SBX read lengths within each bin were calculated and plotted (**Figure 1E**). The fraction of cDNA covered by at least one SBX reads as a function of length was also calculated from the same counts (**figure 1G**).

### Error Rate Comparison

Using aligned reads, error rates were computed both overall and as the per-position incidence of matches, substitutions, insertions, and deletions along the read length. The calculation was performed with a correction for true biological sequence variation and aligner limitations. Specifically, we ignored reads aligning in VDJ genes, HLA genes and MT genes that are either recombinant (leading to incorrect alignments since splice motifs expected around large jumps do not exist) or have high natural variation. To avoid inflating error rate estimates due to background mutation level (overall metrics) or in highly expressed genes (per position), we used two corrections. First, we used DeepVariant to generate a VCF based exclusively on the PacBio reads that was then used in all calculations. Second, we compared PacBio and SBX in tandem, identifying overlapping CB-UMI pairs shared between them that are also classified as FSM, ISM, se_FM or se_IM in PacBio by LRAA Sqanti-like. If a substitution, insertion, or deletion was observed at the same genomic position in both datasets or annotated in the VCF, it was classified as a true mutation or PCR error and counted as a match in both. In rare cases where the same variant appeared as a substitution in one dataset and as a closely spaced insertion-deletion pair in the other, no correction was applied. Given the difference in read sizes between SBX and PacBio, we also limited the calculation to the part of the CB-UMI pairs that were covered by both.

For both SBX and PacBio, error rates were calculated on the raw reads, raw reads in shared barcodes, CUDLL reads, and CUDLL reads in shared barcodes. For SBX, we also calculated them over the set of CUDLLs supported by at least 3 or 10 reads to investigate the effect of read support (**Table S2**).

### Single-cell object construction and integration

Per-platform gene and isoform sparse matrices were loaded into Seurat (v5.4.0) under R (v4.4) as paired RNA and isoform assays and merged into a single object covering 2,995 shared barcodes. For each assay, counts were log-normalized (*NormalizeData*), variable features were selected with the vst method (3,000 features for RNA, 5,000 for isoform), and the resulting feature space was scaled (*ScaleData*) and reduced with PCA (*RunPCA*, npcs = 50). Platform batch effects were corrected with Harmony (*RunHarmony*, group.by.vars = “sample”, 30 dimensions). Independent Harmony embeddings were computed for the RNA and isoform assays. Shared-nearest-neighbour graphs and Louvain clusters (*FindNeighbors*, FindClusters, algorithm = 1) were computed across resolutions 0.4 - 1.2 resolution 0.4 was used for all downstream analyses. UMAP embeddings were generated from each Harmony reduction (*RunUMAP*, 30 dimensions).

### Cell-type annotation

Clusters at resolution 0.4 were manually annotated based on canonical marker expression and FindAllMarkers output, yielding two label tiers used downstream, a fine cell-type label and a coarse RNA lineage.

### VDJ chain calling and CDR3 concordance analysis

VDJ chains were called with MiXCR (*v4*.*7*.*0, preset generic-ont, --species hsa, --rna*) on per-platform alignment CUDLL FASTQs, producing per-cell-barcode clonotype tables and per-clonotype CDR3 summaries. Productive chain calls were retained and joined to the shared single-cell object by cell barcode.

To restrict analyses to authentic receptor expression and exclude chain calls arising from ambient cell-free RNA, a gene expression derived chain-call mask was applied in two stages. First, every chain call was required to originate from a cell barcode present in the platform specific gene expression cell whitelist, removing chain calls captured in empty droplets. Second, the chain class was required to match the cell’s RNA-derived lineage. TCR chains (TRA, TRB, TRG, TRD) were retained only in T-cell-lineage cells and BCR chains (IGH, IGK, IGL) only in B-lineage cells, removing within-cell ambient contamination in which cell-free transcripts of the opposite lineage are co-captured in a real cell’s droplet. Cells annotated as Low QC (higher fraction of ambient transcripts) or as pDCs (known to express BCR transcripts) were excluded from off-target denominators. For the main analysis we applied a permissive per-chain depth threshold of ≥1 supporting UMI and ≥1 supporting CUDLL read (denoted u1/r1); threshold dependence was characterised by repeating the analysis across the full UMI × read grid (u = 1–10, r = 1–20; 200 combinations) per sample–receptor combination.

For each chain in each cell barcode, candidate CDR3 nucleotide sequences were ranked by a local score combining read count, UMI count and within-sample clonotype prevalence; the highest-scoring candidate was assigned as the top call. To resolve cases in which competing candidates of comparable score yielded different top calls between platforms, a pair-exact rescue was applied. For each barcode and chain present on both platforms, if the PB and SBX candidate sets contained at least one nucleotide-identical CDR3 in common, that shared sequence was assigned as the top call on both sides. The rescue rule altered only top-call selection among already-called chains (the called-cell denominator was unchanged by construction). CDR3 concordance was reported as the fraction of barcode chain instances with identical CDR3 nucleotide sequences between PB and SBX, computed both per chain (TRA, TRB, TRG, TRD, IGH, IGK, IGL) and per receptor pair (TRA + TRB; IGH + IGK/L).

### Differential isoform analysis

Differential isoform testing was performed on the same single-cell object after restricting to same-cluster matched cells; barcodes present in both platforms whose RNA cluster assignment was identical between PB and SBX (n = 2,647 of 2,995 shared barcodes, comprising 5,294 cells). For each cluster meeting a minimum size of 20 cells per platform, both gene-level (RNA assay) and isoform-level (isoform assay) cluster-vs-rest differential expression was computed with Seurat FindMarkers (Wilcoxon rank-sum test; logfc.threshold = 0, min.pct = 0.05); markers were called significant at Benjamini–Hochberg-adjusted p < 0.05 with |log2FC| ≥ 0.25. An isoform was classified as “shared”, “PB-only”, or “SBX-only” within each cluster according to whether it reached significance on both platforms, PB only, or SBX only, and these per-cluster status tags were aggregated across cell-type lineages (T-cell, B-cell, NK, Myeloid, Platelet, Low QC) for length and concordance analyses.

### Length-dependent detection bias

To test whether transcript length explains platform-unique differential isoform expression calls, the binary outcome PB-only versus SBX-detected was modelled by logistic regression on log10(transcript length) (glm(is_pb_only ∼ log10_transcript_length, family = binomial)). A natural-spline alternative (ns(log10_transcript_length, df = 4)) was fit as a non-linearity sensitivity check, where AIC was used to compare the two models. Observed PB-only fractions were binned in fixed-width log10 bins and reported with Wilson 95% confidence intervals.

**Table S1:** Read-processing, barcode-correction, alignment, and deduplication metrics for SBX and PacBio sequencing. Read counts and retention rates are shown across barcode filtering, alignment, shared-cell selection, and CUDLL deduplication. Percentages are calculated relative to either raw reads or the preceding processing step, as indicated.

| Processing stage | SBX reads | SBX % of raw | SBX Δ from parent | SBX % of parent | PacBio reads | PacBio % of raw | PacBio Δ from parent | PacBio % of parent |
| --- | --- | --- | --- | --- | --- | --- | --- | --- |
| Raw reads (PacBio = post-Skera) | 9,537,554,778 | 100.00% | — | — | 69,707,639 | 100.00% | — | — |
| After reorienting | — | — | — | — | 69,562,789 | 99.79% | 144,850 | 99.79% |
| After adapters anchoring & trimming | 6,043,640,876 | 63.37% | 3,493,913,902 | 63.37% | 60,069,585 | 86.17% | 9,493,204 | 86.35% |
| Barcoded total (exact + corrected) | 5,983,382,491 | 62.73% | 60,258,385 | 99.00% | 59,966,643 | 86.03% | 102,942 | 99.83% |
| ↳ Barcode exact match | 5,492,622,036 | 57.59% | 551,018,840 | 90.88% | 59,386,631 | 85.19% | 682,954 | 98.86% |
| ↳ Barcode error-corrected | 490,760,455 | 5.15% | 5,552,880,421 | 8.12% | 580,012 | 0.83% | 59,489,573 | 0.97% |
| ↳ Not exact/corrected after trimming (trimmed – exact – corrected) | 60,258,385 | 0.63% | 5,983,382,491 | 1.00% | 102,942 | 0.15% | 59,966,643 | 0.17% |
| Aligned with minimap2 | 5,481,487,661 | 57.47% | 501,894,830 | 91.61% | 59,932,374 | 85.98% | 34,269 | 99.94% |
| ↳ Barcoded but not aligned ((exact + corrected) – aligned) | 501,894,830 | 5.26% | 5,481,487,661 | 8.39% | 34,269 | 0.05% | 59,932,374 | 0.06% |
| In shared valid cell barcodes | 4,609,507,533 | 48.33% | 871,980,128 | 84.09% | 49,142,043 | 70.50% | 10,790,331 | 82.00% |
| Deduped reads that aligned with minimap2 | 47,554,580 | — | 5,433,933,081 | — | 19,988,552 | — | 39,943,822 | — |
| Deduped reads in shared valid cell barcodes | 36,833,140 | — | 10,721,440 | 77.45% | 16,288,489 | — | 3,700,063 | 81.49% |

**Table S2.** Sequence accuracy and error profiles of SBX and PacBio reads before and after CUDLL consensus correction. Match and error rates are shown before and after CUDLL correction. V(D)J, HLA, and mitochondrial loci were excluded, and VCF-adjusted analyses account for shared biological variants and PCR-derived errors.

| Sample/config | Reads | Filtered Reads (w/o VDJ/HLA/MT) | Match rate | Substitution rate | Insertion rate | Deletion Rate | Error rate |
| --- | --- | --- | --- | --- | --- | --- | --- |
| PacBio (VCF adjusted) | 59,932,374 | 55,034,681 | 0.9984 | 0.0004 | 0.0006 | 0.0007 | 0.0016 |
| PacBio shared bc (VCF adjusted) | 49,142,043 | 44,727,148 | 0.9984 | 0.0003 | 0.0006 | 0.0007 | 0.0016 |
| PacBio CUDLL (VCF adjusted) | 19,988,552 | 18,361,087 | 0.9992 | 0.0002 | 0.0003 | 0.0003 | 0.0008 |
| PacBio CUDLL shared bc (VCF adjusted) | 16,288,489 | 14,820,237 | 0.9992 | 0.0002 | 0.0003 | 0.0003 | 0.0008 |
| SBX (VCF adjusted) | 5,481,487,661 | 4,945,125,684 | 0.9901 | 0.0059 | 0.0021 | 0.0019 | 0.0099 |
| SBX shared bc (VCF adjusted) | 4,609,507,533 | 4,124,924,220 | 0.9901 | 0.0059 | 0.0021 | 0.0019 | 0.0099 |
| SBX CUDLL (VCF adjusted) | 47,554,580 | 43,609,705 | 0.9939 | 0.0037 | 0.0013 | 0.0012 | 0.0061 |
| SBX CUDLL shared bc (VCF adjusted) | 36,833,140 | 33,587,717 | 0.9953 | 0.0035 | 0.0012 | 0.0011 | 0.0047 |
| SBX CUDLL shared bc 3copies (VCF adjusted) | 19,796,095 | 17,988,744 | 0.9983 | 0.0013 | 0.0004 | 0.0005 | 0.0017 |
| SBX CUDLL shared bc 10copies (VCF adjusted) | 17,363,308 | 15,807,596 | 0.9986 | 0.0011 | 0.0003 | 0.0004 | 0.0014 |
| PacBio shared bc & UMI, FSM/ISM/se_FM/se_IM UMIs only, shared length (VCF + shared errors adjust) |  | 32,884,524 | 0.9993 | 0.0001 | 0.0002 | 0.0003 | 0.0007 |
| SBX shared bc & UMI, FSM/ISM/se_FM/se_IM UMIs only, shared length (VCF + shared errors adjust) |  | 2,783,441,927 | 0.9906 | 0.0057 | 0.0019 | 0.0017 | 0.0094 |
| PacBio CUDLL shared bc & UMI, FSM/ISM/se_FM/se_IM UMIs only, shared length (VCF + shared errors adjust) |  | 9,956,942 | 0.9997 | 0.0001 | 0.0001 | 0.0002 | 0.0003 |
| SBX CUDLL shared bc & UMI, FSM/ISM/se_FM/se_IM UMIs only, shared length (VCF + shared errors adjust) |  | 9,956,998 | 0.9996 | 0.0002 | 0.0001 | 0.0001 | 0.0004 |

**Table S3:** Concordance between LRAA assignments of PacBio and SBX CUDLLs. Counts are CUDLLs after restricting to the shared barcode whitelist. Assignment categories are mutually exclusive and are evaluated from strict to loose. The top candidate has maximal frac_assigned for a UMI; “good” candidates have frac_assigned >= 50% of the top candidate. Confident assignments only have 1 (shared) assignment.

| Category | Count | % of PacBio | % of SBX | % of common | % of total (union) |
| --- | --- | --- | --- | --- | --- |
| CUDLLs in PacBio | 12,211,648 | 100.00% | — | — | — |
| └ Unique to PacBio | 797,831 | 6.53% | — | — | 3.36% |
| CUDLLs in SBX | 22,970,955 | — | 100.00% | — | — |
| └ Unique to SBX | 11,557,138 | — | 50.31% | — | 48.62% |
| Common CB-UMI (PacBio ∩ SBX) | 11,413,817 | — | — | 100.00% | 48.02% |
| ├ Same confident transcript_id | 7,077,615 | — | — | 62.01% | 29.78% |
| ├ Same confident gene_id | 940,010 | — | — | 8.24% | 3.95% |
| ├ Same top transcript_id | 2,925,166 |  |  | 25.63% | 12.31% |
| ├ Same top gene_id | 230,405 |  |  | 2.02% | 0.97% |
| ├ Overlap in good candidate transcript_id | 3,689 | — | — | 0.03% | 0.02% |
| ├ Overlap in good candidate gene_id | 2,186 | — | — | 0.02% | 0.01% |
| └ No assignment overlap | 234,746 | — | — | 2.06% | 0.99% |

## Author Contributions

C.G. developed CUDLL and implemented the associated read-processing, barcode/UMI correction, and performed benchmarking analyses used to evaluate read-length recovery, splice-structure preservation, and sequence accuracy. G.A-E performed CUDLL downstream single-cell analysis, including differential isoform expression, VDJ/CDR3 recovery and concordance analyses. A.B., A.S., S.L., M.C., and D.A.B. assisted with library preparation and sequencing. S.J.Y. and M.E.K. developed the pipeline for SBX read trimming and filtering, with contributions from H.Y and B.J.H. H.Y., B.J.H., and A.K. contributed to data analysis and interpretation. G. A-E, C.G. and A.M.A wrote the manuscript. V.P., N.H., and N.L. contributed to interpretation and manuscript revision. Roche scientists contributed to SBX sample processing, sequencing, data generation, data processing, analysis, and interpretation. M.K. and A.M.A. conceived and supervised the project.

## Acknowledgements

This work was supported by a collaboration agreement between Roche and the Broad Institute (Broad Clinical Labs). G.A-E. is a Cancer Research Institute Irvington fellow supported by the Cancer Research Institute (CRI5800). A.M.A. is a Merkin Institute Fellow supported by the Broad Institute.

